# Anthropogenic noise can decrease tomato reproductive success by hindering bumblebee-mediated pollination

**DOI:** 10.1101/2024.11.21.624309

**Authors:** Zsófia Varga-Szilay, Gergely Szövényi, Gábor Pozsgai

**Author notes:** Corresponding authors: Zsófia Varga-Szilay, and Gábor Pozsgai.

## Abstract

Anthropogenic noise is a little-studied type of pollution that negatively affects the physiology, nervous function and development of insects, thereby it has the potential to disrupt even key ecological services such as pollination. Here, we investigate the effects of anthropogenic noise on the pollination success of the *Bombus terrestris* on tomatoes in controlled conditions. We expect that bumblebees avoid flowers exposed to noise more than flowers in non-noisy environments, leading to less efficient pollination and lower fruit quality. The experiment was conducted in Hungary, in 2023. Three treatments were applied to randomly chosen flowers: noisy (with played traffic noise); and two non-noisy, one allowing bumblebees and one excluding them. The flowers were enclosed with nets before maturity to avoid pollination, opened exclusively during treatment, and re-enclosed for three more days post-treatment. We recorded the marketing value of the fruits and the number of seeds they produced. We found no significant differences in the marketing value of fruits among treatments, but the number of seeds differed significantly suggesting that anthropogenic noise has a substantial effects on bumblebee-mediated pollination. Although these effects may be mitigated by habituation, loud external noise of various machines (e.g. irrigation systems) within polytunnels is still likely to contribute to the everyday noise exposure of bumblebees and could thus potentially lead to hidden economic losses in production. Therefore, further research is needed to understand the behavioural effects of both direct and indirect noise pollution on bumblebees.

## Introduction

In human-disturbed environments, multiple anthropogenic stressors threaten biodiversity and the healthy functioning of ecosystems (Wagner et al., 2021; Willmott et al., 2022). Among those, anthropogenic noise, as a global phenomenon, is an intense, however less studied type of pollution that has been proven to negatively affect the physiology, nervous function and development of many animal taxa, invertebrates, vertebrates, aquatic and terrestrial species alike (Blickley & Patricelli, 2010; Kight & Swaddle, 2011; Kunc & Schmidt, 2019; Shannon et al., 2016; Slabbekoorn et al., 2018).

Anthropogenic noise can cause alterations in physiological responses, such as heightened stress levels, increased pain, or elevated stress hormone production (Barber et al., 2010; Kight & Swaddle, 2011; Shannon et al., 2016). These physiological effects often lead to increased fitness costs, potentially reducing reproductive success and survival. For instance, studies in birds suggest that adult growth can be impaired and longevity be reduced even when eggs are exposed to traffic noise (Meillère et al., 2024). Moreover, noise disrupts the perception of environmental cues and social interactions of animals that rely on acoustic signals (Brumm & Slabbekoorn, 2005; Classen-Rodríguez et al., 2021; Siemers & Schaub, 2010) (e.g. mating interactions), significantly affecting not only birds (Engel et al., 2024; Halfwerk & Slabbekoorn, 2013) and mammals (Slabbekoorn & Peet, 2003), but also invertebrates, such as molluscs (e.g. Gigot et al., 2024; Serdar et al., 2024) and insects (e.g. Orci et al., 2016; Welsh et al., 2023). The effects of noise pollution are not confined to acoustically oriented animals but extend to organisms with no direct links to the acoustic realm (Senzaki et al., 2020). For instance, highway noise induces physiological stress in monarch caterpillars (Davis et al., 2018) and even dragonflies without acoustic receptors inhabiting environments adjacent to noisy areas are negatively impacted (Senzaki et al., 2020).

To mitigate the effects of noise pollution, animals may either avoid noisy environments or develop compensatory strategies. For instance, some species modify their acoustic signals by increasing the amplitude (Lombard effect) (Brumm & Todt, 2002; Katti & Warren, 2004; Nemeth & Brumm, 2010) or shifting frequencies to higher ranges in noisy environments (Gil & Brumm, 2014; Lampe et al., 2012; Nemeth et al., 2013; Shieh et al., 2012). Others alter their activity patterns, or, ultimately, they might move to quieter places (Brumm, 2013; Warren et al., 2006). However, both compensation and avoidance can lead to significant behavioural changes, such as altering in movement, habitat use (Duarte et al., 2011), foraging patterns and foraging efficiency (e.g. Luo et al., 2015; Siemers & Schaub, 2010), or communication patterns (e.g. Brumm & Slabbekoorn, 2005; Naguib, 2013). Nonetheless, these behavioural shifts come with energy costs, which, similarly to physiological effects, can negatively affect survival and reproductive success (Kight & Swaddle, 2011). However, in some cases, like in crickets, plasticity can, at least partially, buffer against these fitness costs if noise exposure is constant (Welsh et al., 2023).

Behavioural changes, on the other hand, can also have cascading effects at the community level (Barber et al., 2010; Senzaki et al., 2020) and can indirectly influence species interactions (e.g. predator–prey interactions, (Barton et al., 2018), through which they can hamper key ecosystem services such as pollination (Francis et al., 2012; Phillips et al., 2021). For instance, Phillips et al. (2021) found that while road noise does not affect the density of bumblebees (*Bombus* spp.), the number of flower visits and the time bees spend on flowers are significantly higher at sites farther from roads. Additionally, noise pollution may impact vibration-based buzz pollination and short-range acoustic communication between individuals, further influencing pollination success (Bunkley et al., 2017). Thus, noise ultimately disrupts the effectiveness of pollination and can therefore lower the reproductive success of both wild plants and crops (Guenat & Dallimer, 2023).

Since the 1980s, commercial bumblebees (e.g., *Bombus terrestris audax* in Europe) have been widely used in both glasshouse and open-field cultivation (De Ruijter, 1997; Velthuis & Doorn, 2006), as they are among the most efficient pollinators for a wide variety of popular crops like tomatoes (Cooley & Vallejo-Marín, 2021; De Luca & Vallejo-Marín, 2013), eggplants (Mondal et al., 2022), and strawberries (Gudowska et al., 2024). Even crops that are capable of self-fertilization are well-known to produce larger, heavier, and better quality fruits, such as sweet peppers (Roldán Serrano & Guerra-Sanz, 2006), when bumblebees transport the pollen to the stigma. One key advantage of bumblebees in pollinating plants over other bees, such as honey bees, lies in their ability to use specific frequency vibration of their thoracic muscles to shake out the pollen from the anthers of some plants (termed ‘buzz pollination‘) (De Luca & Vallejo-Marín, 2013; Vallejo-Marín, 2021). Therefore, these crops are likely to be disproportionately impacted by noise-induced decline in pollination activity.

Despite the well-documented impacts of anthropogenic noise pollution on birds (e.g. Halfwerk & Slabbekoorn, 2013) and marine mammals (e.g. Hastie et al., 2021; Sørensen et al., 2023) there remains a significant knowledge gap regarding to invertebrates (Guenat & Dallimer, 2023; Jerem & Mathews, 2021; Morley et al., 2014; Shannon et al., 2016; Sordello et al., 2020), and particularly on the effects of noise on pollinator behaviour. Previous experiments have focused on the effect of acoustic vibrations of bumblebees on pollination (e.g. Vallejo-Marín, 2021; Woodrow et al., 2024), but how pollination success is altered in noisy environments has not yet been examined. As a result, our understanding on how noise affects these vital pollinators of cultivated plants remains incomplete. Additionally, it is still unknown to what extent bumblebees avoid highly noise-polluted areas, and it remains uncertain whether noise pollution impacts crop pollination success and yield quality.

Thus, we investigated the effects of anthropogenic noise on the pollination success of the *Bombus terrestris* (Linnaeus, 1758) and fruit production on tomatoes (*Solanum lycopersicum* L., Solanales: Solanaceae) in a polytunnel setting in Hungary. We used tomatoes as they are a key crop, and although they are self-compatible, high yields and high-quality tomato fruits require buzz pollination (Picken, 1984). They are also the best-studied buzz-pollinated plant species (Cooley & Vallejo-Marín, 2021). Since we expect that bumblebees spend less time on flowers exposed to noise, or avoid them completely, we predict less efficient pollination and lower fruit quality in noisy environments than in non-noisy ones. In line with this, we formulated the following hypotheses:

**H1:** Exposure to traffic noise results in a decrease in the number of tomato seeds, which is a reliable measure of pollination success and an important indicator of the plant’s reproductive success.

**H2:** Traffic noise negatively impacts the market value of fruits, thereby affecting their saleability and, consequently, their economic value for the producer.

## Material and methods

### Study sites

The experiment was conducted in Szentes (46°39’09.2” N 20°13’58.4” E), located in the Southern Great Plain region of Hungary (**Fig. 1**), under controlled conditions within a polytunnel, between May 18 and 28, 2023. The polytunnel measured 10 x 70 x 5 m, covering an area of 700 m², and was heated with thermal water. The cultivation followed integrated pest management (IPM) practices and the Aruba tomato variety was grown. The tomato plants, which were planted in September of the previous year, were approximately three meters tall and at the 25th truss stage during the experiment. Ten rows of plants were arranged in the polytunnel, and within these rows, a total of twelve bumblebee nests were placed. Exclusively *Bombus terrestris* was used in the polytunnel, with the nests positioned at its centre. The experiment was conducted in the first two and the fourth rows from the right side of the polytunnel. In the first row, 25 out of 139 plants were selected, with 32 flowers chosen for the study. In the second row, 14 plants from a total of 148 were included, and 17 flowers were selected. For the third row, 8 plants out of 127 were involved, with 9 flowers chosen. The plants and flowers within each set were randomly selected and spaced as far apart as possible to ensure treatment independence.

**Fig. 1:**
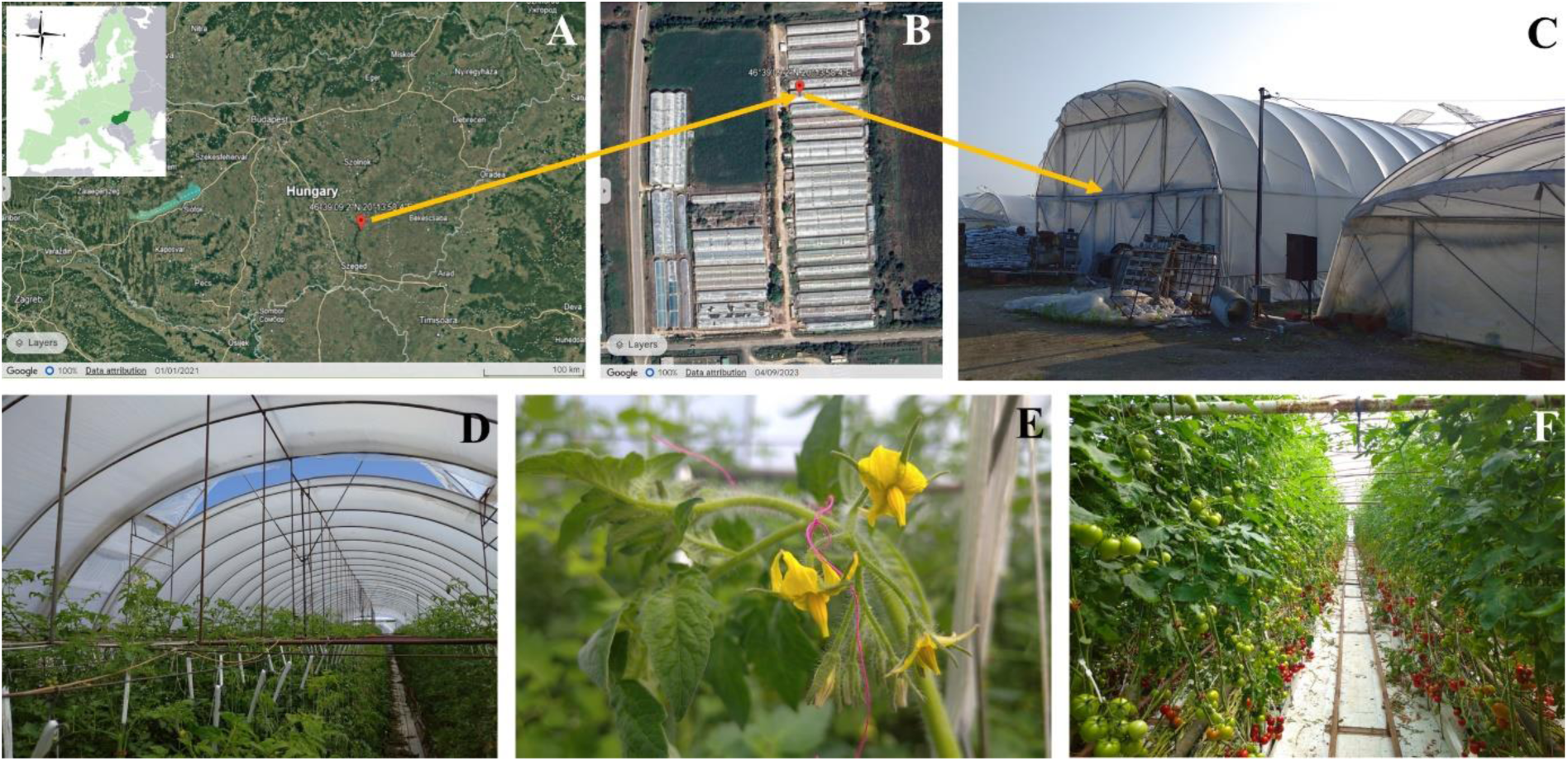
The study site location in Szentes, Hungary (A), the aerial view and the exterior (B, C), and the interior of the polytunnel where the experiment took place (D-F).

### Experimental design

The experiments were conducted between 7:00 and 18:00 each day. The selected tomato flowers were numbered and enclosed with nets before maturity (n = 62), released exclusively during treatment, and then re-enclosed for three more days (**Fig. 2**).

**Fig. 2:**
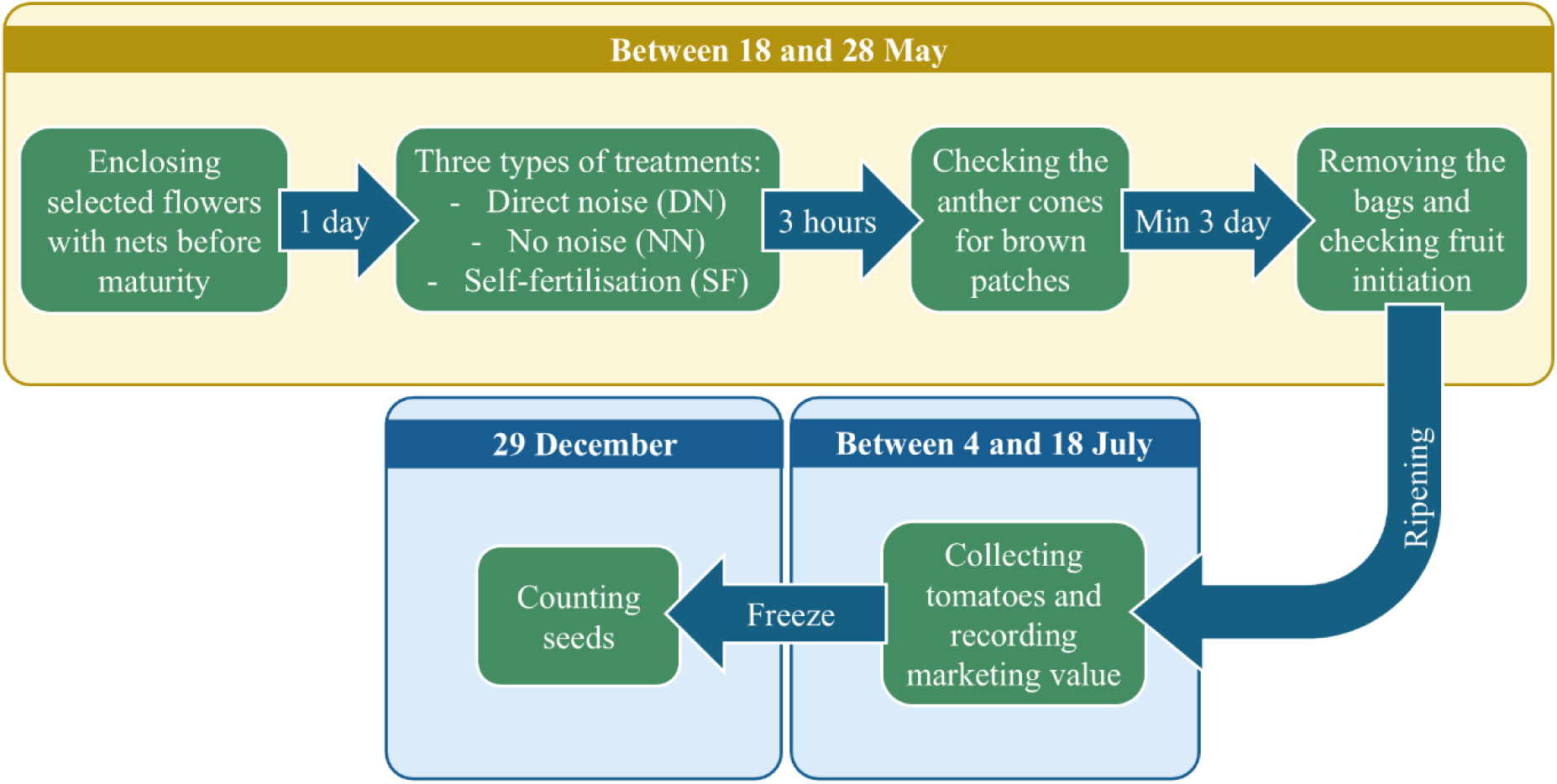
The experimental and data collection process. The yellow frame indicates the onsite processes and data collection, while the blue frames indicate post-experiment data collection.

We used 20 repetitions with three types of treatments on selected flowers: a noisy (DN) with played traffic noise (hereafter noise) from a speaker (see below, **Fig. 3A**), a non-noisy (NN) with a paper dummy speaker (**Fig. 3B**) matching in size and colour yet not producing sound, and treatment with the complete exclusion of bumblebees (SF, **Fig. 3C**) thereby ensuring self-fertilisation of the flower, without any played noise or dummy speaker. Both the real speaker and the dummy were placed directly next to the selected flowers, at a maximum distance of 10 cm (see **Fig. 3A, B**). For both noisy and non-noisy treatments, the bags were removed for 3 hours, for the time of treatment. In the noisy treatment after 5 minutes of the pre-treatment period, the flowers were exposed to noise. No noise was played for the non-noisy and self-fertilisation treatments.

**Fig. 3:**
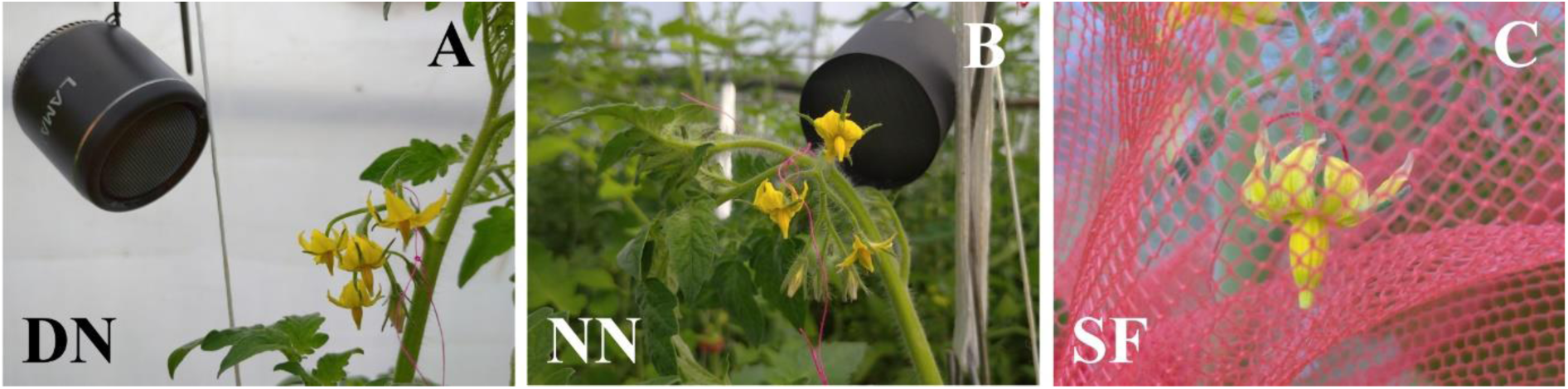
The three types of the study treatment on tomato flowers: (A) noisy (DN); (B) non-noisy (NN); and (C) the complete exclusion of bumblebees (self-fertilisation, SF).

Environmental conditions, including temperature and humidity, were not modified for the experiment and remained as typically found within the polytunnel, optimised for tomato growth. The temperature at the beginning of the experiments averaged 25.79°C (± 3.12°C), while at the end of the experiments, the average temperature was 28.09°C (± 2.96°C). The background noise was recorded at the start of each experiment day. For this, we used a VOLTCRAFT SL-10 sound level meter with a sensitivity of 0.1 dB, a frequency range of 31.5 Hz to 8 kHz, and a sound level measurement range of 30 to 130 dB(A). The average of the lowest background noise levels was 43.03 dBA (± 3.73 dBA SD), and the average of the highest of that was 60.93 dBA (± 4.93 dBA SD). The traffic noise (the audio source: https://www.zapsplat.com) was played using a LAMAX Sphere2 Mini Bluetooth 5.1 speaker with a frequency range of 150 Hz to 20 kHz. The average of the lowest treatment noise levels was 69.96 dBA (± 10.76 dBA SD), and that of the highest was 85.96 dBA (± 7.11 dBA SD).

### Data collection

We recorded the total number of flowers on each cluster and noted the position of the flower subjected to the treatment. Immediately after the treatments, we assessed whether pollination had happened by carefully observing the anther cones (**Fig. 4A**) on noisy and non-noisy treatments (it was not relevant for the self-fertilisation treatment) and checking for the presence of bite marks (brown discolouration) on the flowers. After a minimum of three days (since fertilisation occurs around 24-50 hours post-pollination), we permanently removed the bags and recorded whether there was any sign of berry initiation (**Fig. 4B**).

**Fig. 4:**
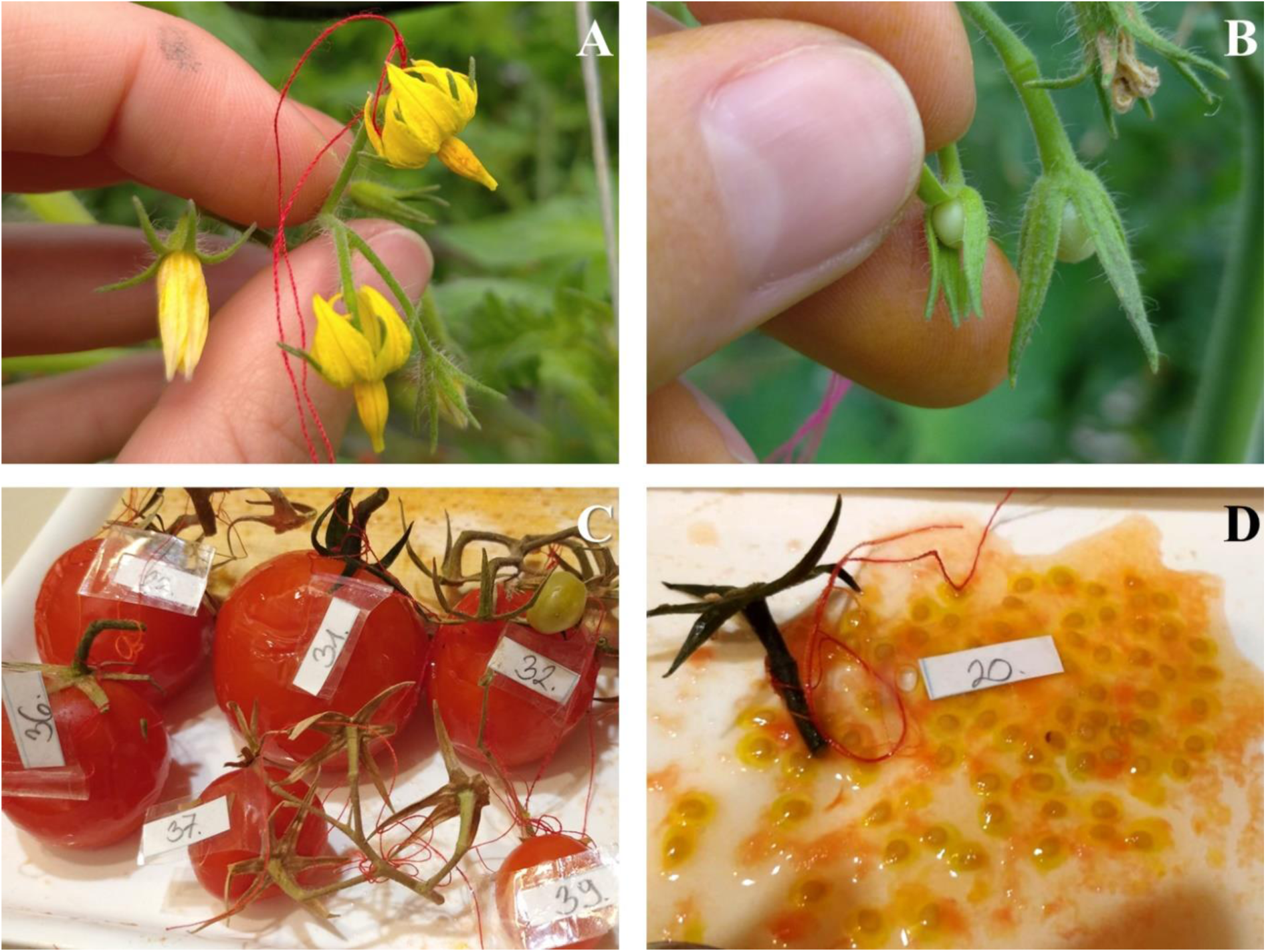
Data collection by observing the anther cones (A), checking the sign of berry initiation (expanding fruit, B), recording the marketing value (C), and counting the total number of seeds of tomato fruits (D).

After fruit ripening, we collected the tomatoes or made a note if the numbered tags were missing (**Fig. 4C**). The fruits were classified for their marketing value by the producers who were unaware of the applied treatments. Very small, green or lowest value fruits were scored to Category 1, those which were sold in bulk as Category 2, and those which were sold in assortments of seven and six were scored to Category 3 and Category 4 (highest values), respectively. Non-developed fruits were scored as Category 0. After classification, the collected tomatoes were frozen for subsequent seed counting (**Fig. 4D**).

### Statistical analysis

We performed a two-sample proportion test to compare whether there is a difference between the noisy (DN) and non-noisy (NN) treatments based on brown patches. Additionally, Pearson’s Chi-squared test was used to assess variations among the treatments regarding berry initiation (expanding fruit).

To analyse the marketing value of the fruit by treatment, first, we converted marketing value categories to numerical variables and then we used a generalised linear model (GLM) with a Poisson distribution to appropriately handle the integer data. For the analysis of the number of seeds by treatment, we also utilised a generalised linear model (GLM) with a Poisson distribution. Flowers that failed to set fruit were excluded from this latter analysis. We used estimated marginal means (EMMs) from the “*emmeans*” R package (Lenth et al., 2024) for pairwise comparisons to examine the differences between treatments. We ensured that the variances were homogeneous in all occasions with the Goldfeld-Quandt test, and tested the normality with the Shapiro-Wilk test when the seed numbers were compared. The data analysis and visualization of the results were conducted in the R (R Core Team, 2021).

## Results

Of the flowers included in the experiments (n = 62), we recorded on 41 flowers (18 and 23 flowers of noisy and non-noisy treatments, respectively) whether brown patches were present and from all flowers if the initial fruit was developed. Of the 62 flowers 44 developed fruits, 2 plants with 2 marked flowers died before developing fruits, 4 flowers failed to set fruit, while the outcome of 12 flowers is unknown.

### Brown patches and fruit initiation

The two-sample proportion tests comparing noisy (DN, n = 18) and non-noisy (NN, n = 23) treatments indicated no significant difference between the treatment groups the presence of brown patches (χ² < 0.001, p = 1.00, 95% CI = -0.28 to 0.33). Moreover, Pearson’s Chi-squared test indicated no significant difference in initial fruit development among all treatments (χ² = 0.556, df = 2, p = 0.757).

### Marketing value of the fruit

The marketing value of the fruit, on average, tended to be higher in the non-noisy (NN) treatment than in the other two groups (**Fig. 5**), yet the differences were not significant (analysis of deviance, deviance = 33.327, df = 45, p-value = 0.767). The pairwise comparisons did not show significant differences among the treatments (**Table 1**).

**Fig. 5:**
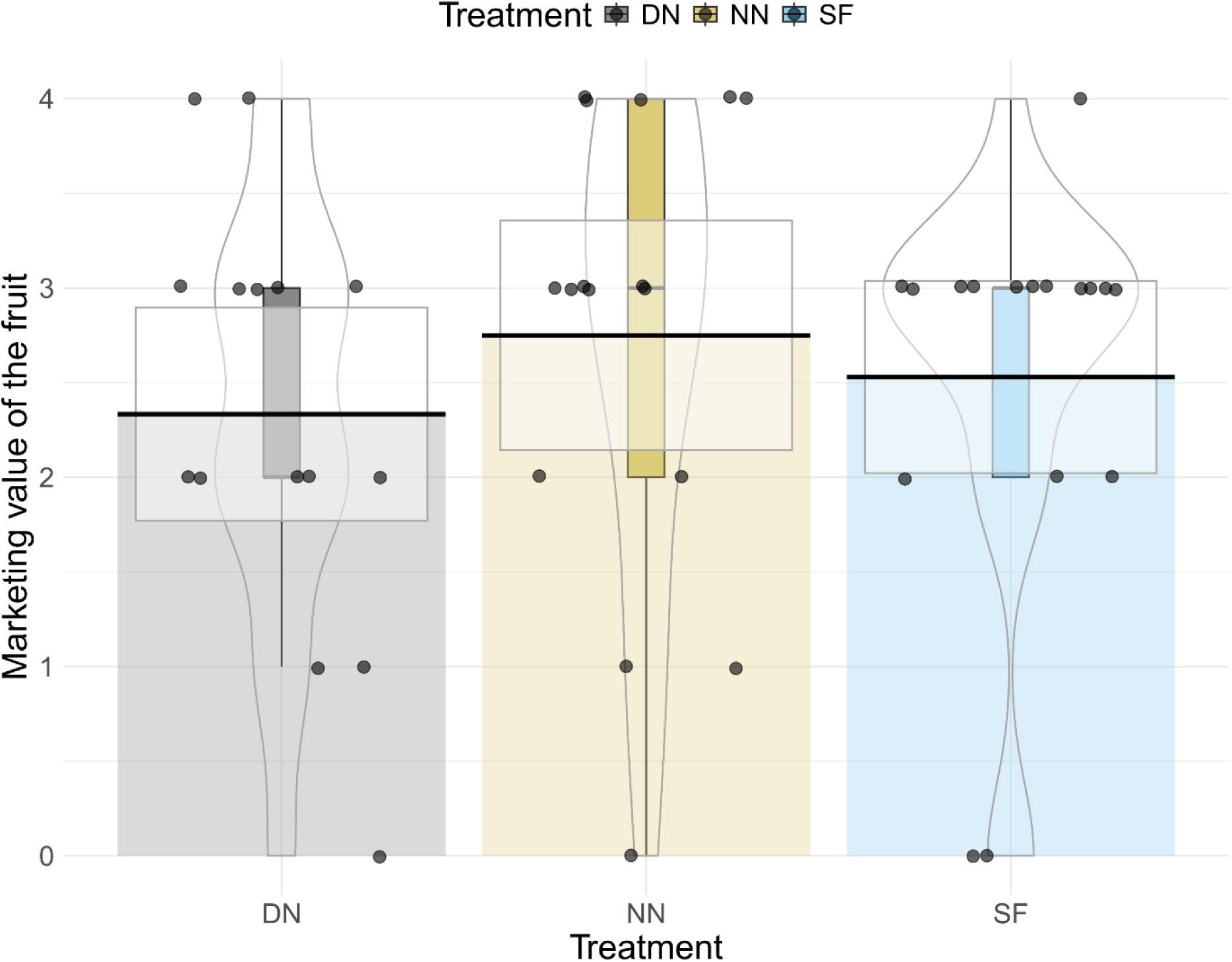
The marketing value of the fruit by the three types of treatments, noisy (DN, n = 15), non-noisy (NN, n = 16), and self-fertilisation (SF, n = 17). Dots represent the category of the marketing value of the fruit of individual tomato samples. The categories are defined as follows: 0 – undeveloped fruit, 1 – very small or green fruit, 2 – fruit sold in bulk, 3 – fruit sold in assortments of seven, and 4 – fruit sold in assortments of six. Box plots display the median and interquartile range, while violins show density, mean, and credible intervals.

**Table 1:** Comparisons of marketing values of tomatoes between treatment pairs with estimated marginal means (no p-value adjustment). Treatments: DN: noisy (n = 15), NN: non-noisy (n = 16), SF: self-fertilisation (n = 17).

| Treatments | Estimate | SE | df | z ratio | p-value |
| --- | --- | --- | --- | --- | --- |
| DN – NN | -0.164 | 0.226 | Inf | -0.725 | 0.4682 |
| DN – SF | -0.081 | 0.228 | Inf | -0.354 | 0.7230 |
| NN – SF | 0.084 | 0.214 | Inf | 0.390 | 0.6966 |

### Number of seeds

The generalised linear model implied significant differences among treatments in seed numbers (analysis of deviance, df = 41, deviance = 897.61, p < 0.001). Differences were also significant between each treatment pair (**Table 2**), with fruits under the non-noisy (NN) treatment producing the most seeds, and those under noise (DN) showing the lowest seed counts (**Fig. 6**).

**Fig. 6:**
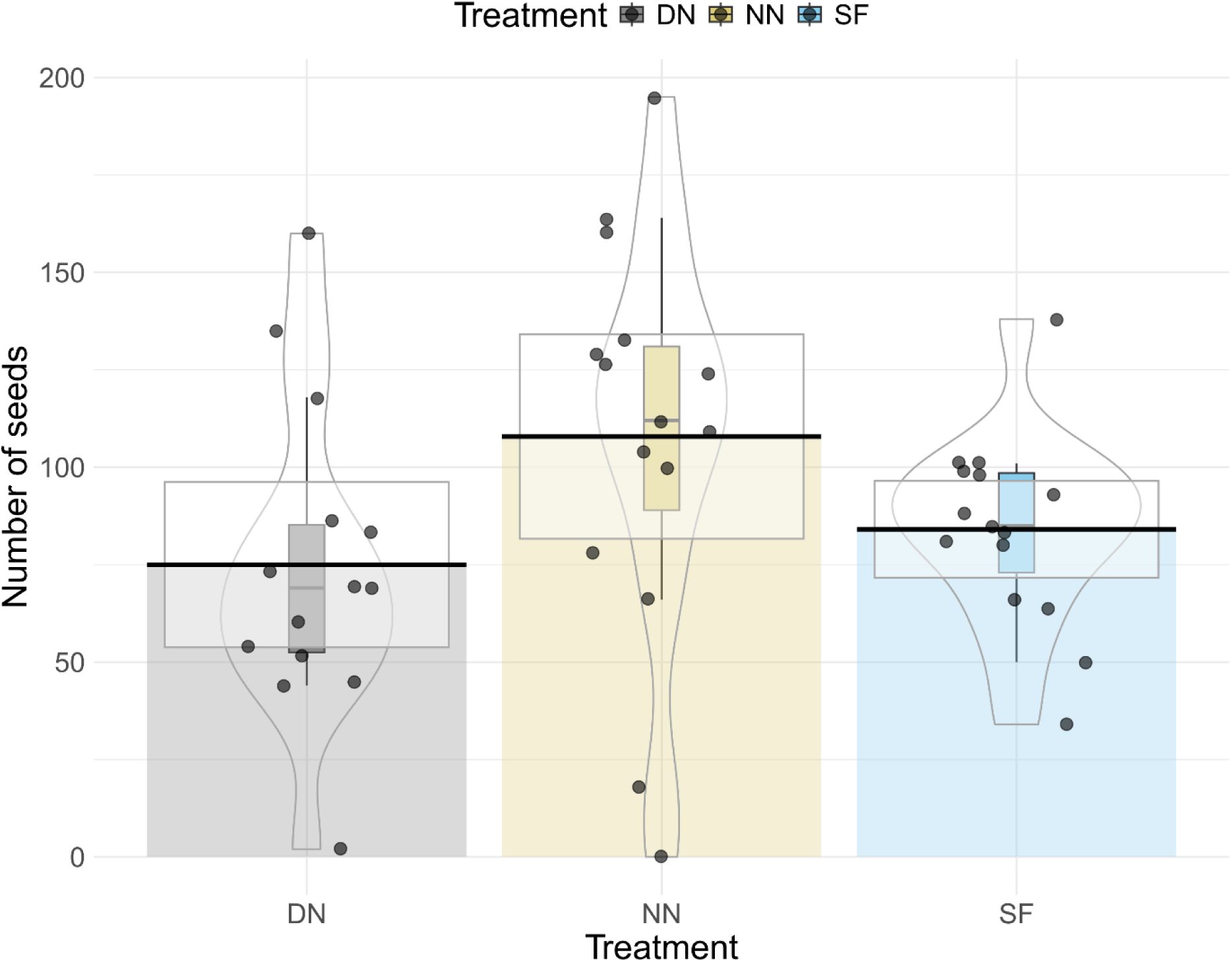
The number of tomato seeds separated by treatments, noisy (DN, n = 14), non-noisy (NN, n = 15), and self-fertilization (SF, n = 15). Dots represent the seed number of individual tomato samples. Box plots display the median and interquartile range, while violins show density, mean, and credible intervals.

**Table 2:**
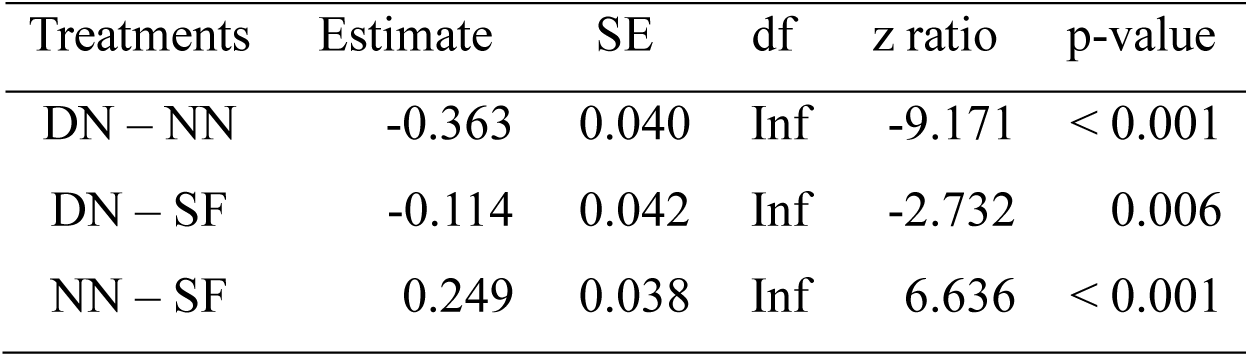
Comparisons of the number of seeds between treatment pairs with estimated marginal means (no p-value adjustment). Treatment: DN: noisy (n = 14), NN: non-noisy (n = 15), SF: self-fertilisation (n = 15).

| Treatments | Estimate | SE | df | z ratio | p-value |
| --- | --- | --- | --- | --- | --- |
| DN – NN | -0.363 | 0.040 | Inf | -9.171 | < 0.001 |
| DN – SF | -0.114 | 0.042 | Inf | -2.732 | 0.006 |
| NN – SF | 0.249 | 0.038 | Inf | 6.636 | < 0.001 |

## Discussion

In this study, we examined how anthropogenic noise affects the pollination success of *Bombus terrestris* on tomatoes under semi-controlled polytunnel conditions. Our findings confirm that traffic noise has a measurable significant negative effect on fruit quality, measured as seed production (**H1** holds), suggesting an adverse impact on bumblebee pollination behaviour. However, the significantly lower seed count in noisy treatment than in the self-fertilisation one also implies a negative impact on the plants which are difficult to set apart from those on pollinators. On the other hand, noise did not influence the marketing value of the fruits (**H2** was not supported). This lack of significant difference in marketing value among treatments suggests that the direct economic impacts of anthropogenic noise may be more nuanced than initially anticipated, at least at the coarse level we used for value characterisation. However, the consistent and relatively loud background noise within the polytunnel where the experiment was conducted may also have masked the differences between our treatments. Moreover, the fact that fruits under noisy treatments produced less seeds than self-fertilising flowers suggests a direct negative effect of noise on fruit quality and allows the speculation that consumer value, in some unmeasured parameters important for human consumption, such as vitamin and nutrient content, may also be negatively affected. Additionally, for other crop types grown outside of polytunnels/greenhouses, seed count decreasing could have important implications for future yield.

Indeed, within greenhouses and polytunnels, daily operations produce a variety of noises from sources such as automated irrigation systems, and human activity, including tasks and conversations, or radios playing music from early morning until the end of the workday. Since continuous noise exposure is likely to accumulate over time (Kok et al., 2023), common noise sources – beyond traffic or industrial activities – could also affect bumblebee-mediated pollination. These, in turn, can potentially lead to substantial disruptions in flower-visitation activity and pollination success and can have broader implications for yield quantity and quality. Moreover, these continuous background noises may also negatively affect brood size and quality in bumblebee nests, which may have a measurable economic cost in bumblebee production facilities.

On the other hand, behavioural flexibility, or the ability to exhibit plastic responses (e.g. bird song plasticity, Slabbekoorn, 2013) to stable but intrusive sounds, could help buffer some effects of noise, allowing animals to function (survive and reproduce) in high-noise urbanised or agricultural environments. Studies on invertebrates (e.g. grasshoppers, Lampe et al., 2014) also suggest that the capacity to ‘tune out’ uniform stimuli may help maintain essential behaviours even under anthropogenic noise pollution. Thus, in polytunnels, the continuous exposure to repeated stimuli of background noise may lead bumblebees to habituate to these sounds, reducing immediate stress responses. Yet, this adaptation may only apply when noise remains relatively constant, as more variable noise is likely to impose higher physiological and behavioural costs (Naguib, 2013). Thus, non-constant noise of machinery or radio is still likely to affect bumblebees’ behaviour and diminish pollination efficiency.

Beyond its impact on foraging behaviour, research on field crickets and scallops has shown that larval or juvenile exposure to noise can disrupt the normal development of invertebrates, potentially impairing adult behaviours related to survival and reproduction (de Soto et al., 2013; Welsh et al., 2023). While similar effects in bumblebee larvae are currently speculative, such impacts could potentially affect colony success rates and reproductive output, highlighting an area open for further investigation.

Indeed, it is plausible to expect that noise pollution also influences various stages of bumblebees’ lifecycle, including the hibernation ability of queens and the reproductive success of colonies, such as larval development rates and the number of reproductive offspring. However, developmental noise exposure may also contribute to habituation and noise tolerance in adult bumblebees. Nevertheless, if managed bumblebee colonies are raised in noisy environments which reduces their sensitivity to noise and potentially limits their observable behavioural responses, they still undergo early-life stress that may negatively influence growth and can have oblique deleterious effects on physiology which ultimately decrease pollination efficiency. This further implies that commercial bumblebee producers could benefit from controlling the environmental noise in their facilities.

### Limitations

Our sample size was constrained by the setup of the polytunnel, where only three out of the ten rows had plants lowered to a height (still nearly 3 metres) that allowed access to the flowers. Though we examined the anther cones for the presence of bite marks as an indicator of flower visitation immediately after treatments, in fact, it can take up to four hours for these brown marks to become fully visible. Therefore, the presence or absence of these marks immediately after treatment could not be considered a reliable indicator for flower visitation, thus it must be interpreted with caution. Lastly, we did not directly measure bumblebee activity during the treatment, which may have provided additional insights into the impact of noise on their foraging behaviour.

### Conclusion and future perspectives

With this study, we laid the foundations of how bumblebee-mediated pollination efficiency is impacted by noise. Even in our relatively small-scale study, we found deleterious negative effects of noise on tomato pollination. This, from an economic perspective, underlines that noise pollution can diminish pollination efficiency which could lead to unforeseen losses in crop yields, particularly near to urban infrastructure. Our study underscores the importance of identifying and managing noise sources within agricultural systems, such as polytunnels and greenhouses, where bumblebees are necessary for high-quality production. Yet, currently, there is a lack of information on how anthropogenic noise affects entire bumblebee colonies, creating a significant gap in our understanding. Addressing this is essential, as understanding these potential effects is crucial for developing strategies to mitigate noise-related disruptions and to ensure the sustainability of pollination services. Further studies are needed to understand how different types of anthropogenic noise can alter the behaviour of both wild and commercially used pollinators, and consequently, pollination efficiency. To fully elucidate the nuanced impacts of noise, future experiments should involve a broad variety of crops (e.g. strawberry or pepper), and other commercially used bumblebee species (e.g. *Bombus impatiens*).

Moreover, efforts to utilise acoustic signals to estimate bumblebee activity (e.g. (Miller-Struttmann et al., 2017)) and the use of artificial-sound-assisted pollination for crops like tomatoes or strawberries that respond to sonication to replicate natural pollination (Dingley et al., 2022) are in the rise. However environmental noise may disrupt the necessary signals and hamper the further development of these methods. Thus, focusing research on the effects of noise on pollination is inevitable from this perspective as well.

Ultimately, gaining a comprehensive understanding of the ecological effects of noise pollution on bumblebees’ and other pollinators’ behaviour has a high potential for developing practices that better support pollinator health and well-being, ensuring the sustainability of pollination-dependent agriculture in a rapidly urbanising world.

## Author Contributions

Conceptualization, Z.V.S., G.S. and G.P.; Methodology, Z.V.S. and G.P.; Software, Z.V.S. and G.P.; Validation, Z.V.S. and G.P.; Formal Analysis, Z.V.S. and G.P.; Investigation, Z.V.S.; Resources, Z.V.S. and G.P.; Data Curation, Z.V.S. and G.P.; Writing – Original Draft Preparation, Z.V.S. and G.P.; Writing – Review & Editing, Z.V.S., G.S. and G.P.; Visualization, Z.V.S. and G.P.; Supervision, G.P. and G.S.; Project Administration, Z.V.S.; Funding Acquisition, G.P. All authors have read and agreed to the published version of the manuscript.

## Data Availability

The underlying computer code is available in the GitHub repository https://github.com/zsvargaszilay/buzz_amidst_noise/

## Acknowledgements

We are grateful to Gábor Deák (Szentes-Bio Kft.) for choosing the experimental site and for helping me with the organization throughout the experiment. We also thank the Szentesi Pa-ri Kft. for providing the experimental site and for contributing to the follow-up data collection. Last but not least, we thank the Talent Support Council of ELTE Eötvös Loránd University for supporting the experiment.

